# A modified Cas9 scaffold allows extension of the virus-induced gene editing technology to the large P*otyvirus* genus

**DOI:** 10.64898/2026.02.02.703200

**Authors:** Fernando Merwaiss, Verónica Aragonés, Arcadio García, José-Antonio Daròs

## Abstract

Plant viruses are recognized as rapid and effective vectors to deliver CRISPR-Cas reaction components into plants, a strategy termed virus-induced gene editing (VIGE). However, VIGE is limited by the host range of the viral vectors. Development of new viral vectors to target a broad range of plant species will potentially enable the delivery of the editing components to new cultivars. Potyviruses (genus *Potyvirus*) comprises the largest group of plant RNA viruses. The main limitation of potyviral vectors to express a non-coding RNA consists of potential insertion of stop codons that interrupt the large open reading frame that encompass most potyviral genome. This is the case with the *Streptococcus pyogenes* Cas9 sgRNA scaffold, which contains stop codons in all three possible frames. In this work, we first built on a visual reporter system targeting the two homeologs of *Nicotiana benthamiana Magnesium chelatase subunit I* (*CHLI*). Second, we developed a tobacco etch virus (genus *Potyvirus*)-derived vector for VIGE by engineering a modified Cas9 scaffold, free of stop codons, to maintain the potyviral polyprotein reading frame while ensuring effective editing. This vector self-replicates and moves systemically, delivering sgRNAs efficiently throughout the plant. This allowed to obtain plants exhibiting a white phenotype with their four alleles edited through *in vitro* regeneration from infected leaves, and also to produce edited progeny. We further demonstrated the vector utility in tomato. Given the conserved biological properties within the genus *Potyvirus*, these findings must be broadly applicable to other potyviruses, expanding the reach of the VIGE technology.

## Introduction

The advent of targeted genome editing technologies, particularly those derived from the CRISPR-Cas systems, has revolutionized plant science and offers immense promise for crop improvement and fundamental genomic studies^1,2^. These technologies enable precise DNA modifications that can confer beneficial traits, such as enhanced tolerance to biotic and abiotic stresses or increased nutrient values^3^. However, the widespread application of genome editing in plants is often constrained by the requirement for labor-intensive and genotype-specific *in vitro* tissue culture, typically involving *Agrobacterium tumefaciens*-mediated transformation or biolistics for delivering gene-editing reaction components, typically Cas nucleases or derivatives and one or more guide RNAs^4,5^. These conventional methods can be time-consuming, lead to unintended genome changes or damage, and pose regulatory restrictions due to the integration of exogenous DNA in the host genome.

To overcome this bottleneck, virus-induced gene editing (VIGE) has emerged as a promising alternative for delivering CRISPR components directly into living plants^6–9^. VIGE systems leverage plant viruses as transient delivery vehicles, allowing efficient expression and systemic spread of the CRISPR-Cas reaction components. Among the advantages, VIGE bypasses or significantly reduces the need for *in vitro* tissue culture, accelerates the editing process, enables editing at the whole plant level, and, critically, if using an RNA viral vector, facilitates the production of transgene-free gene-edited plants in a DNA-free process.

Both RNA and DNA viruses have been engineered as VIGE vectors, each with distinct advantages and applications. RNA viral vectors are particularly appealing because their replication occurs in the host cytoplasm with no DNA intermediates, typically ensuring that the resultant edited plants are free from foreign DNA, thus avoiding regulatory issues associated with transgenes^1,7^. However, a major challenge for many RNA viruses is their limited cargo capacity, which makes delivering large Cas nucleases, such as the frequently used *Streptococcus pyogenes* Cas9 (SpCas9, ∼4.1 kb), problematic^10–13^. For this reason, many VIGE systems utilize RNA viruses to deliver only the single-guide RNA (sgRNAs) into Cas9-expressing plants^4,14,15^, although recently efforts are being done to incorporate Cas nuclease expression into VIGE system to achieve true transgene-free processes^11,13,16–18^.

One of the most widely used RNA VIGE vectors is tobacco rattle virus (TRV). This bipartite virus, which consists of two positive-sense single-stranded genomic RNAs, exhibits a broad host range, infecting over 400 plant species^1,13,15^. TRV-based vectors have been successfully used to deliver sgRNAs into transgenic plants constitutively expressing SpCas9^4,15^. A crucial advance for improving heritable genome editing with TRV involved the augmentation of the sgRNAs features by fusing RNA mobility-promoting sequences like a fragment of the *Flowering locus T* (*FT*) mRNA or some tRNAs^4,19–21^. These modifications enable sgRNAs to move into the germline cells of the meristems, which is essential for transmitting mutations to the next generation. For example, the *FT*-modified sgRNAs significantly increased heritable editing frequencies in *N. benthamiana*, with some progeny displaying complete gene knockout phenotypes. Interestingly, even with these modifications the TRV vector remains with low seed-transmissibility, leading to a virus-free edited progeny^7^. Potato virus X (PVX), a flexible, filamentous plus-strand RNA virus, is another effective vector, especially for species in the family *Solanaceae* ^6,22,23^. Its structure minimizes limitations on insert size, making it suitable for delivering larger genes like Cas9 alongside sgRNAs^6^. PVX vectors have achieved high editing efficiencies in *N. benthamiana*. Importantly, PVX has been successfully employed for multiplex editing using unspaced sgRNA arrays and has enabled heritable VIGE in tomato^22–24^. Interestingly, PVX was found to induce biallelic edited progeny even without explicit RNA mobility signals^25^.

Despite significant advancements, VIGE still faces challenges, such as the cargo limitation that difficult expression of large CRISPR-Cas systems^10,12^, or the exclusion from meristematic germline cells that limits heritability^11^. Host range is another factor that limits VIGE applicability, as each viral vector only infects a particular set of plant species. Development of new VIGE vectors to widen host range is required to reach new crop species and cultivars. In this regard, genus *Potyvirus* comprises the largest group of RNA viruses infecting plants^26^. However, potyviral gene expression strategy based on production of large polyproteins pose difficulties to express non-coding RNAs, such as SpCas9 sgRNAs. In this work, we aimed to incorporate potyviruses into the VIGE toolbox. To achieve this, we first used more classic TRV and PVX vectors to adapt the SpCas9 sgRNA scaffold to the stop-codon-free format required for potyviral expression. Second, we confirmed efficient genome editing in SpCas9-expressing *N. benthamiana* using different potyviral vectors, opening the VIGE technology to this large genus of RNA viruses that collectively infect a broad range of host plants.

## Results

### Harnessing a visual reporter system to evaluate VIGE in plants

As a first step to expand the array of VIGE vectors in plants, we considered that a reliable visual marker to specifically detect edited tissues was crucial. Therefore, our first goal was to establish a robust model for evaluating the *in vivo* editing efficiencies of various viral vectors in a *N. benthamiana* SpCas9-expressing plant. Drawing on existing literature, we selected the *Magnesium chelatase subunit I* (*CHLI*) as an ideal visual marker. *CHLI* is involved in the chlorophyll biosynthetic pathway and, consequently, in developing the characteristic green color of plants. Notably, because *N. benthamiana* is an allotetraploid, two homeologous loci for this gene exist on chromosomes 14 and 12, which we named here *CHLI-A* and *CHLI-B*, respectively. Using the CRISPOR computational tool^27^, we designed three different sgRNAs (named sgRNA-1, 2 and 3) potentially able to simultaneously target exon 3 of both *CHLI* homeologs, aiming for good predicted efficiency while minimizing potential off-targets (**Figure S1a**).

To empirically test these sgRNAs, we cloned them into TRV and PVX vectors, whose utility in VIGE has already been validated. The 5’ *FT* mobile domain was also added to the 3’ end of the sgRNAs to promote heritable genome editing (**Figure 1a**). All six constructs, along with their respective empty-vector versions, were agroinoculated into *N. benthamiana* Cas9-expressing plants. In contrast to mock-inoculated plants, all plants agroinoculated with the viral vectors expressing *CHLI*-specific sgRNAs started to exhibit white sectors in systemic tissue. At 25 days post-inoculation (dpi), samples from upper leaves displaying a white phenotype were collected, and genomic DNA was purified for analysis. Specific primers were designed to selectively amplify fragments of *CHLI-A* or *CHLI-B*, and the PCR products were sequenced. The inference of CRISPR edits (ICE) for each homeolog obtained for both viral vectors assayed was calculated using the EditCo Bio software. A representative time-course picture series showing extension of bleached tissue in the case of TRV-sgRNA1 is presented in **Figure S1b**. See also the **video in Supporting Information** encompassing four weeks after the agroinoculation.

**Figure 1.**
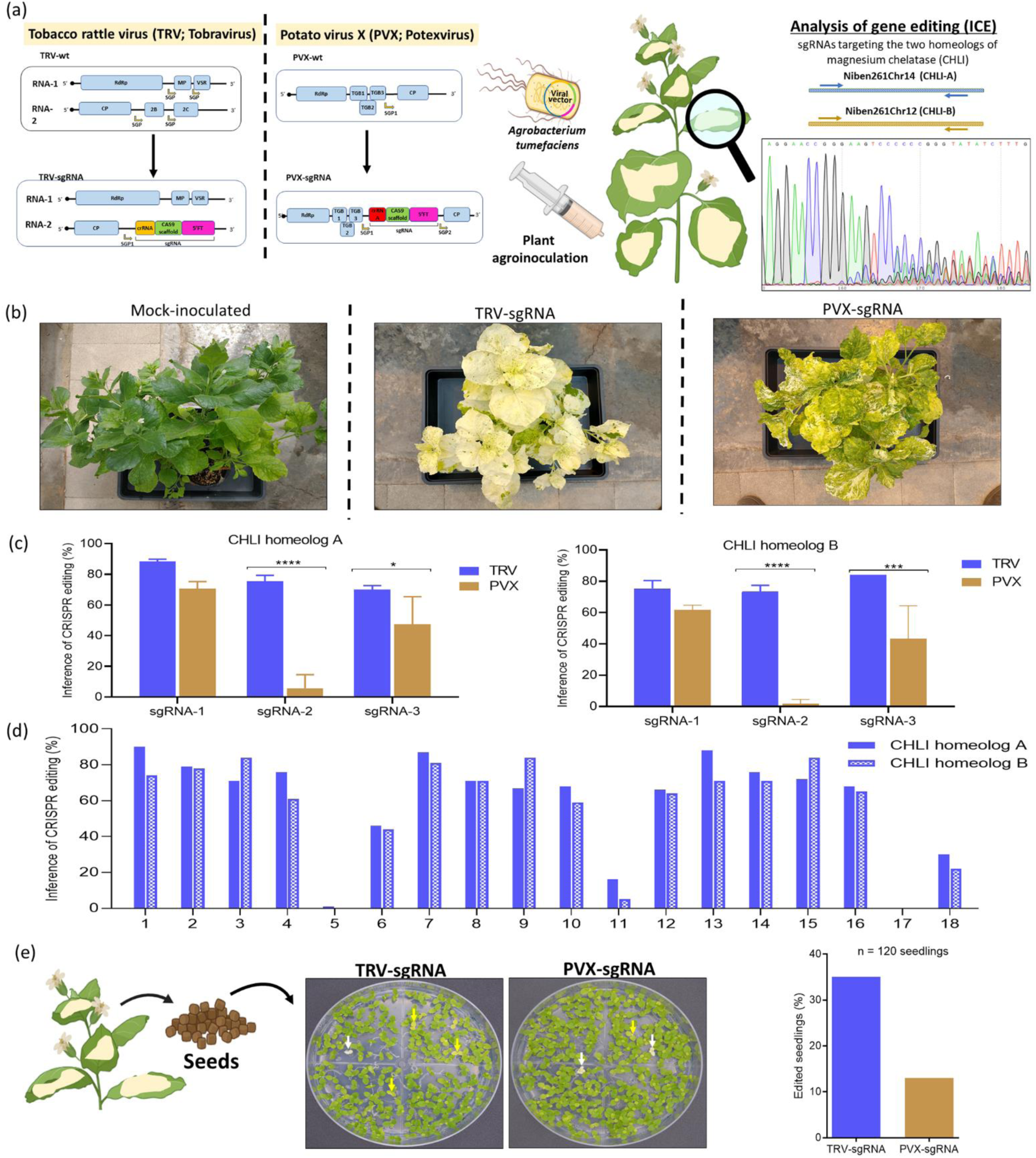
Comparison of the editing efficiency for three different sgRNAs in two different viral vectors. a) The sgRNAs were cloned into two different viral vectors, TRV (left) and PVX (right) in order to empirically compare both, the editing efficiency of the specific sequences and the viral vectors for VIGE. *N. benthamiana* plants were agroinoculated with the different viral vector/sgRNA combinations. At 21 dpi, upper leaves presenting a bleached phenotype were harvested and genome editing was analyzed. b) Representative adult plants mock-inoculated or agroinoculated with TRV or PVX carrying sgRNAs against *CHLI* c) Comparison of the editing efficiency of each sgRNA with both viral vectors for each *CHLI* homeolog (homeolog A, left; homeolog B, right). Statistical analysis was performed using the GraphPad Prism 8 software. Data were analyzed from 3 biological replicates by two-way ANOVA with Tukey’s posttest (*,P<0.05; **,P<0.01; ***,P<0.001; ****,P<0.0001). d) Comparison of the editing efficiency for both homeologs in each analyzed sample. e) Seeds of plants agroinoculated with TRV-sgRNA-1 and PVX-sgRNA-1 were collected and germinated *in vitro* on agar plates supplemented with sucrose. DNA was extracted from 120 seedlings pooled in groups of 3 and the proportion of edited alleles was calculated by sequencing and ICE analysis. White arrows highlight the presence of white seedlings while yellow arrows show yellow seedlings.

Pictures of whole representative plants were taken at 30 dpi. Although all combinations of sgRNAs and viral vectors induced white phenotype patches in the infected plants, the degree of photobleaching was much more homogeneous with the TRV than with PVX (**Figure 1b and S1c**). The ICE results for each homeolog were compared across guides and vectors. Consistent with phenotypic observations, the TRV vector showed significantly higher activity, especially with sgRNA-2 and sgRNA-3, compared to the PVX vector (**Figure 1c**). We also found that sgRNA-1 and sgRNA-3 were more efficient than sgRNA-2 with both vectors, and that this effect was more pronounced for the PVX vector. As a complementary analysis, we also represented the data for each individual sample obtained for both viral vectors assayed by graphing the ICE of both homeologs (**Figure 1d**). These results showed that both homeologs were always similarly edited, indicating that in *N. benthamiana* sgRNAs common to both homeologs of CHLI target both genes with same efficacy. Editing patterns obtained were similar for all sgRNAs in both viral vectors, with a predominant presence of +1 base pair (bp) insertions and -1 bp deletions, followed by small deletions up to -8 bp.

To confirm that the observed white phenotypes resulted from an alteration of the chlorophyll biosynthesis pathway, and furthermore, that these results correlated with the ICE values, we measured the amount of chlorophyll in three representative leaves of each infected plant (**Figure S1d**). The chlorophyll levels were inversely correlated to the degree of editing, confirming that knocking out *CHLI* leads to a decrease in chlorophyll accumulation that explains the bleached phenotype. Taken together, these results support that *CHLI* is an excellent marker to visually follow VIGE in *N. benthamiana*. Results also support that the outcome of VIGE approaches in plants depend both on the selected sgRNA and the viral vector used for expression.

### Analysis of heritable VIGE using the *CHLI* reporter system

Building on the results described above, sgRNA-1 was selected for further research. The next step was to evaluate both the differences between TRV and PVX in reaching the germline and the resulting phenotype of edited seedlings. To achieve this, seeds from several flowers of three different infected plants (for each vector) were collected and germinated *in vitro* on agar plates supplemented with sucrose (**Figure 1e**). DNA was extracted from 90 seedlings, and the proportion of edited alleles was calculated by sequencing fragments of both homeologous genes and performing the previously described ICE analysis (**Figure 1e**). The results showed that although both vectors produced seeds with at least one edited allele, the efficiency with TRV was approximately three times higher than PVX (35% *vs.* 13%).

In several independent experiments of seed germination, we observed that seedling phenotypes could be categorized into three distinct groups, green, light green/yellow, and white (**Figure S2a**, green, yellow and white arrows). We then sought to establish a precise correlation between these phenotypes and the number of edited alleles, which in the case of *N. benthamiana* could range from 0 in wild-type plants to 4 in plants with both loci completely edited. For this, the sequence of both homeologs was analyzed in 96 seedlings (32 of each phenotype). For each analyzed homeolog, ICE values obtained were classified in three categories: values near 100% (both edited alleles), values near 50% (one edited allele) and values close to 0% (no edited alleles). Individual seedlings were plotted, indicating their phenotype and the number of edited alleles (**Figure S2b**). A strong correlation was observed. All green seedlings had 0, 1, or 2 edited alleles, while 94% of yellow plants had 3 edited alleles, and 94% of white plants had all 4 alleles affected. Notably, this effect was independent of whether the edited alleles corresponded to *CHLI-A* or *CHLI-B*, supporting a redundant and equivalent function of both genes in *N. benthamiana*.

While white plants lacking functional *CHLI* alleles can only grow on sucrose-supplemented agar plates, yellow plants (with 3 out of 4 edited alleles) can grow in regular soil and produce fertile flowers (**Figure S2c**). We then analyzed how the mutation was transmitted to the progeny. Seeds from a yellow plant were harvested and germinated *in vitro*. Consistent with Mendelian inheritance, the seedlings from these plants exhibited the expected 1:2:1 (white:yellow:green) phenotypic distribution (**Figure S2d**). Taken together, these results indicate that, in the allotetraploid *N. benthamiana*, CRISPR knocking of the *CHLI* marker gene can be easily predicted from the leaf color in both the infected tissues and the progeny plants.

### Design of a modified sgRNA suitable for VIGE using potyviral vectors

After harnessing the *CHLI* visual reporter system for VIGE in *N. benthamiana* in both the infected plant and the progeny, we used this marker to extend the VIGE technology to the large genus *Potyvirus*. In potyviruses, the region between the nuclear inclusion *b* (Nib) and coat protein (CP) cistrons can accommodate exogenous sequences without affecting viral replication, particularly if they are flanked by cleavage sites that allow excision of the extra protein from the viral polyprotein by means of the viral NIa protease (NIaPro). We hypothesized that, even though sgRNAs do not encode functional proteins, we could insert them into this position in the viral genome, provided translation reading frame is conserved and no in-frame stop codons are introduced^28^. Therefore, a tobacco etch virus (TEV)-derived vector carrying sgRNA-1 between the NIb and CP cistrons was designed. NIb/CP NIaPro cleavage site was complemented at both sides of the insert, including synonymous mutations to avoid sequence repetitions (**Figure 2a**). However, SpCas9 scaffold contains stop codons that would prematurely terminate the polyprotein translation in all three frames. To overcome this inconvenience, we designed a modified scaffold with no in-frame stop codons. Building on the work by Nishimasu et al., who examined the impact of different mutations in the SpCas9 scaffold on editing efficiency, we incorporated specific mutations and deletions to create a modified stop-codon-free SpCas9 scaffold, designated here sgRNAm (C25U, G46A and ΔC30-G39) (**Figure 2b**). To test the functionality of this modified scaffold, we first cloned it into the TRV vector (TRV-sgRNAm-1) and compared its editing efficiency with the original scaffold (TRV-sgRNA-1). Agroinoculated *N. benthamiana* plants showed a systemic white phenotype at 30 dpi, which was similar for both vectors (**Figure 2c**). Further, ICE analysis of both *CHLI* homeologs in these plants confirmed that the editing efficiency was not affected by the mutations in the scaffold (**Figure 2d**).

**Figure 2.**
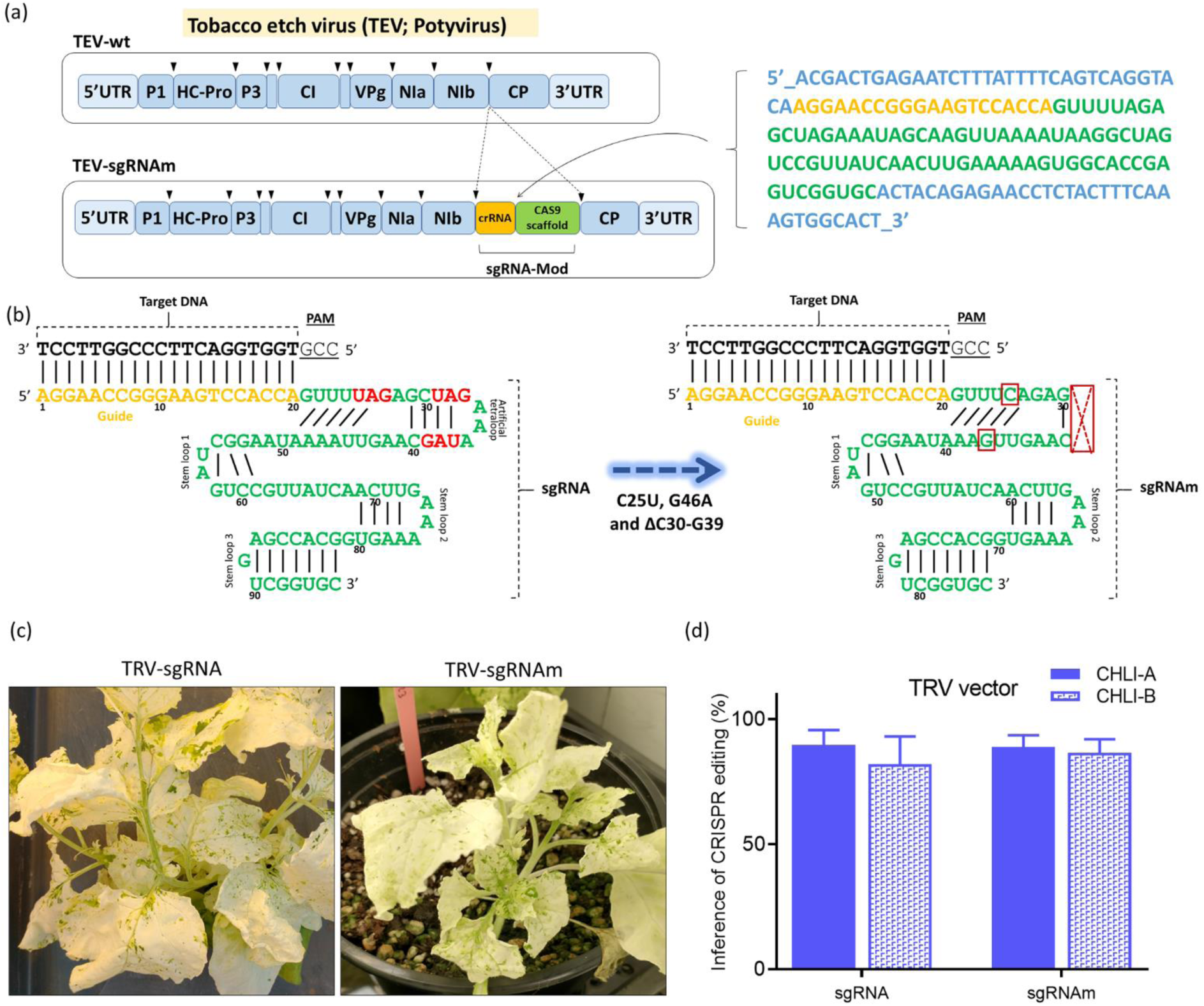
Development of a modified sgRNA suitable for potyviral expression. a) A TEV derived vector carrying the sgRNA between the NIb and CP cistrons was designed. A second TEV-protease recognition site, with a modified synonym nucleotide sequence, was added after the inserted sequence in order to allow cleavage of the exogenous peptide from the polyprotein. b) Given that the sgRNA scaffold contains stop codons (shown in red) that interrupt the large open reading frame in all three possible frames (left), we designed a modified scaffold (sgRNAm) free of stop codons that maintain the polyprotein reading frame (right). c) In order to test the functionality of the modified scaffold, we cloned it in the gold-standard TRV vector (TRV-sgRNAm) and compared the editing efficiency with the original scaffold (TRV-sgRNA). 30 days post agroinoculation, *N. benthamiana* plants presented a systemic white phenotype similar for both vectors. d) ICE analysis of both *CHLI* homeologs confirmed that the editing efficiency was not affected by the modified scaffold.

Next, we transitioned to the TEV vector, where we cloned the sgRNAm-1. We agroinoculated *N. benthamiana* Cas9-expressing plants with both TRV-sgRNAm-1 and TEV-sgRNAm-1 to compare their VIGE efficiencies. White patches appeared in both groups of plants, as representative images taken at 14 dpi showed. However, the TEV-infected plants also presented necrotic symptoms, likely impacting vector functionality (**Figure 3a**). In line with this observation, ICE analysis of both homeologs from the agroinoculated plants showed that TEV induced gene editing in both *CHLI* alleles, albeit at a lower rate than TRV (**Figure 3b**). Unfortunately, TEV-infected plants exhibited limited growth and died after 30 dpi. Given our particular interest in targeting plant germline cells for heritable genome editing, we decided to attenuate the TEV vector to reduce infection symptoms and allow plants to reach the flowering stage^29^. Based on previous results, we designed three different single-point mutations in the HC-Pro cistron, known to attenuate symptoms. These mutations were named AS20a, CLA2, and AS13a^30^. All three attenuated TEV-sgRNAm-1 variants retained their editing activity, but only the CLA2 and AS13a allowed plant survival and seed production in the infected plants (**Figure 3c** and **d**). We observed a positive correlation between the severity of infection symptoms and the efficiency of gene editing, with the TEV-AS13a vector producing the mildest symptoms, but also resulting in the least effective editing. Taken together, these results indicate that VIGE can be opened to the large genus *Potyvirus*, provided modified Cas scaffolds are used that maintain the viral vector translation open reading frame.

**Figure 3.**
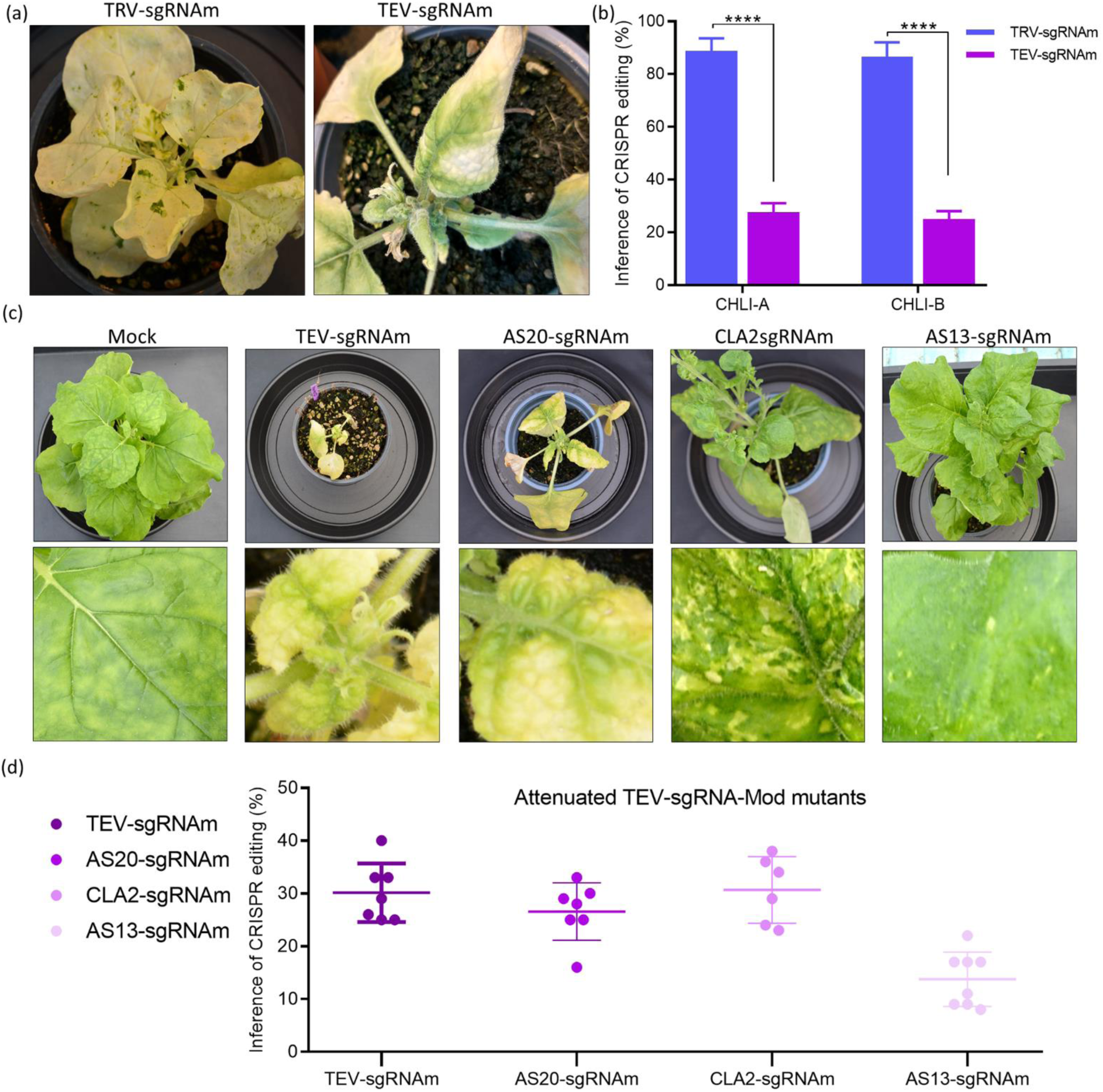
Development of an attenuated potyviral vector for VIGE. a) VIGE with the sgRNAm-1 against CHLI was compared using the TRV and the TEV vectors. Pictures of representative plants were taken at 14 dpi. Plants developed a white phenotype in both cases, but TEV-infected plants presented necrotic symptoms. b) The ICE analysis of the agroinoculated plants showed that TEV is able to induce gene editing in both *CHLI* alleles, although at a lower rate than TRV. Statistical analysis was performed using the GraphPad Prism 8 software. Data were analyzed from 5 biological replicates by two-way ANOVA with Tukey’s posttest (****,P<0.0001). c) Plants agroinoculated with three different TEV vectors developed for VIGE carrying single-point mutations that attenuates its symptoms. d) ICE analysis obtained with the different TEV attenuated vectors. All attenuated TEV variants retained their editing activity while only CLA2 and AS13a variants allowed survival and development of seeds in the infected plants.

### *In vitro* regeneration of edited plants and heritable gene editing using the TEV vector

A major goal of CRISPR technology in plant is to develop varieties with desired gene edits. To achieve this, a strategy relies on *in vitro* regeneration of genetically modified individuals from edited tissue. To assess that edited plants can be regenerated from tissues infected with the TEV vector, we analyzed plants infected with the attenuated TEV-sgRNAm-1 CLA2 and AS13a vectors. After 30 dpi, *N. benthamiana* leaves showed white spots, suggesting the presence of edited tissue. These leaves were harvested and leaf pieces subjected to plant regeneration *in vitro* (**Figure 4a**). Several white plants were obtained, presumably indicating that all *CHLI* alleles were edited. ICE analysis of both *CHLI* homeologs in five plants per condition confirmed that most of them were fully edited. **Figure 4b** illustrates the proportion of edited (white) versus non-edited (green) alleles in regenerated plants. RT-PCR diagnosis of TEV confirmed the presence of the virus in all regenerated plants, as expected.

**Figure 4.**
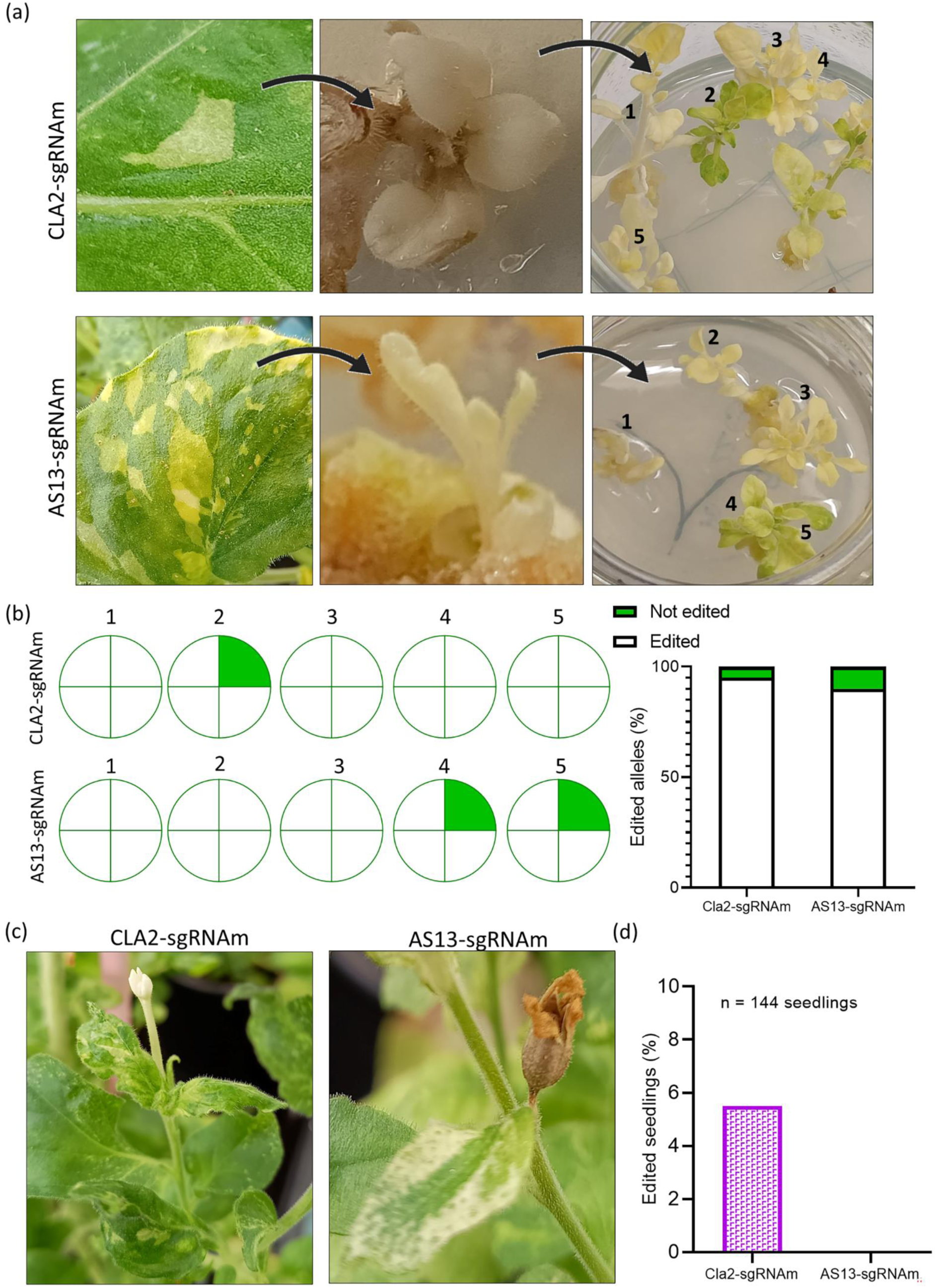
In vitro regeneration of edited plants from TEV-agroinoculated tissue and heritable gene editing. a) Plant leaves with white spots, inferring edited tissues, were collected and in vitro regenerated in agar plates supplemented with sucrose. Several white plants were obtained for both viral vectors. b) ICE analyzes of 5 plants obtained from each condition showed that most of them were completely edited plants. Proportion of edited (white) and non-edited (green) alleles were represented in individual plants (left) or in the complete population assayed (right). c) Representative pictures of flowers near leaves presenting white edited phenotype. d) Seeds from plants agroinoculated with the CLA2 and the AS13a TEV attenuated vectors were germinated in vitro and the presence of edited alleles in both homeologs was evaluated by ICE.

More importantly, we investigated the heritability of CRISPR edits arising directly from plants inoculated with the attenuated TEV-sgRNAm-1 vectors. Based on heterogeneous distribution of viral vectors in infected plants, we collected seeds from flowers adjacent to white tissues (**Figure 4c**). We germinated *in vitro* 72 seeds per vector and used ICE to evaluate edited alleles in both homeologs of the seedlings (**Figure 4d**). Results showed that 5% of seedlings from TEV-CLA2 infected plants carried at least one edited allele. In contrast, no edited seedlings were found in the progeny from TEV-AS13a infected plants. Notably, in contrast to what occurs with the regenerated plants, TEV was not detectable by RT-PCR in the edited progeny. Taken together, these results indicate that, using a TEV vector, edited plants can be obtained both from *in vitro* regeneration of infected tissues or through the progeny of infected plants when using the attenuated TEV mutant CLA2.

### Expanding the use of potyviral vectors for VIGE

Finally, we evaluated whether the TEV vector could be used in species other than *N. benthamiana*, specifically in tomato, cultivar MicroTom. Since TEV symptoms are less severe in tomato MicroTom compared to *N. benthamiana*, we used the wild-type (non-attenuated) TEV-sgRNAm-1 vector for this assay (whose sequence carried one single mismatch against the *CHLI* from tomato). SpCas9-expressing tomato MicroTom plants were agroinoculated and, at 30 dpi, samples were collected from leaves showing symptoms of infection. Although no white patches were observed in these tissues, ICE analysis revealed that nearly 75% of the analyzed samples showed a detectable editing signal, with an average of 10% (**Figure 5a**). A representative Sanger sequencing profile is shown. No editing was detected in the progeny of these edited plants.

**Figure 5.**
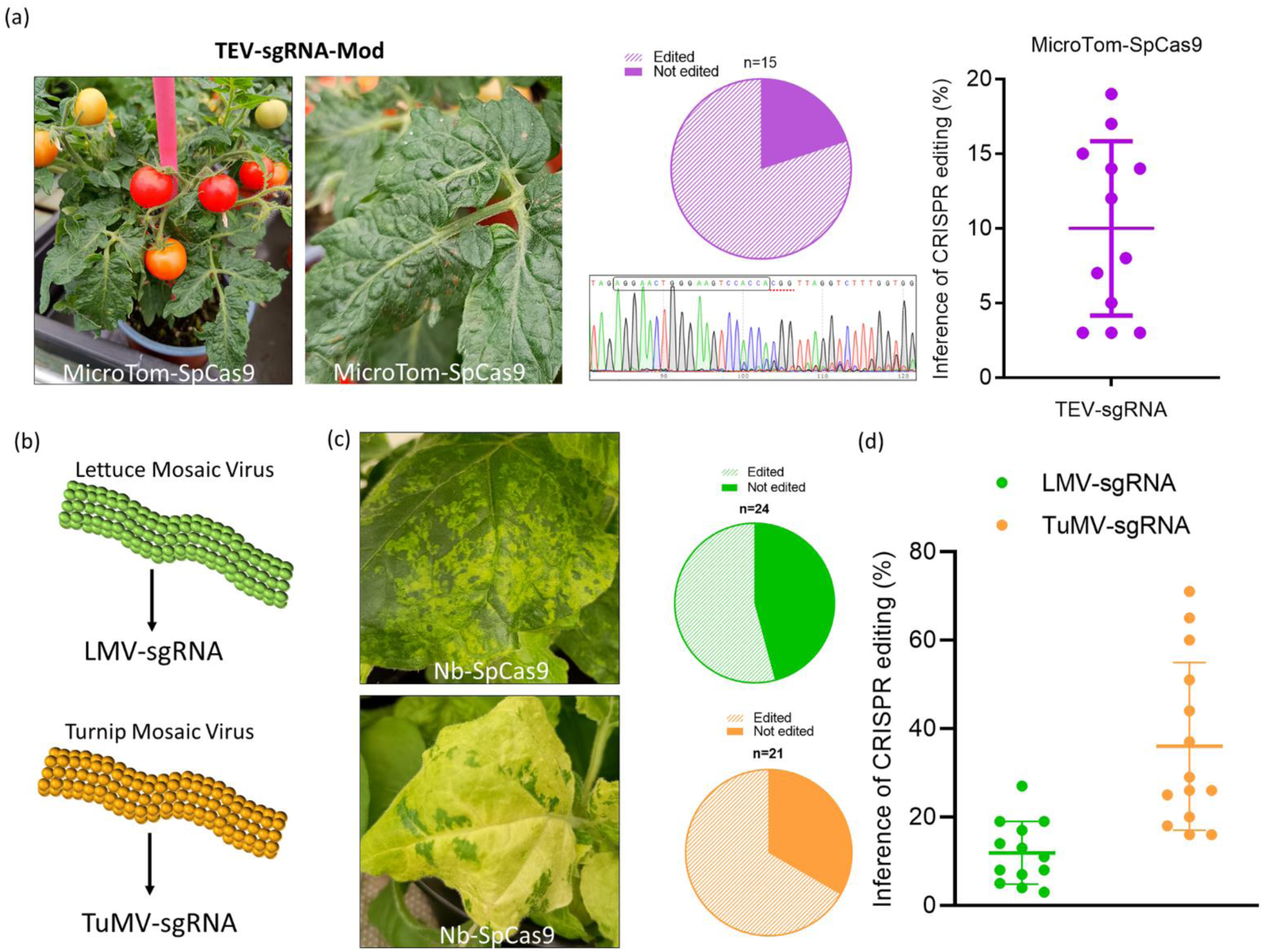
Expanding the use potyviral vectors for VIGE. a) The use of the TEV vector for VIGE was also evaluated in a species of agronomic interest as Micro-Tom tomato expressing Cas9. Pie chart shows the proportion of samples with detectable editing signal, which were then plotted on the ICE graph on the right. A representative Sanger sequencing profile is shown. b) Lettuce mosaic virus (LMV) and turnip mosaic virus (TuMV), whose main host are lettuce and brassica respectively, were designed cloning the same sgRNAm-1 in the same position that for TEV. c) Cas9-expressing *N. benthamiana* plants were agroinoculated with both vectors and tissue samples were collected at 30 dpi. Pie charts represent the proportion of samples that had detectable editing, according to the ICE analysis performed. d) Positive edited samples were plotted.

To ensure that the engineered sgRNAm scaffold was not target specific, we analyzed the editing efficiency of a second marker gene using our potyvirus vector. We chose to target the red fluorescent protein dsRED that is transgenically expressed in the SpCas9-expressing tomato plants. We first designed three different guides following the same procedure than for *CHLI* (R1, R2 and R3), and compared their target efficacy using a TRV vector in *N. benthamiana* plants. Based on these results, we selected the sgRNAm-R1 as the most efficient guide. Tomato plants were then agroinoculated with TEV-sgRNAm-R1. Leaf fluorescence was analyzed 21 dpi with a stereomicroscope. Imaging of one representative leaf of a not infected plant showed full expression of dsRED, compared with two leaves of an agroinoculated plant showing a reduced patched expression (**Figure S5a**). Notably, all the 11 samples analyzed showed positive ICE values (ranging from 8 to 60%), which were plotted on the graph shown (**Figure S5b**). A representative Sanger sequencing profile is included. These results suggest that the usability of the modified sgRNA scaffold designed in this work is not restricted to the *CHLI* marker.

Due to conserved biological properties within the large genus *Potyvirus*, we investigated whether these findings can be directly applied to other potyvirus species, thereby potentially opening the host range of species amenable to this technology. We built vectors derived from lettuce mosaic virus (LMV) and turnip mosaic virus (TuMV) expressing sgRNAm-1 into the identical position as for TEV (**Figure 5b**). We agroinoculated SpCas9-expressing *N. benthamiana* plants to evaluate the vectors efficacy for VIGE. Tissue samples, some showing white patches, were collected at 30 dpi, and ICE analysis of *CHLI-A* was performed. As shown in the pie charts representing the proportion of samples with detectable genome editing, a significant portion of the analyzed samples were successfully edited (**Figure 5c**). Positive edited samples were plotted based on their ICE value, revealing that TuMV produced higher editing efficiency than LMV, likely related to its higher compatibility with the assayed host (**Figure 5d**). Taken together, these results showed that different potyvirus species can be used as VIGE vectors in different host species.

## Discussion

CRISPR-Cas systems have emerged in the last decade as powerful tools for precision genome editing and gene expression control. This technology is particularly attractive to crop science due to the potential straightforward generation of more productive and nutritive cultivars, resistant to pests and pathogens, and better adapted to changing environmental conditions^31^. A successful CRISPR-Cas gene editing application greatly depends on an efficient delivery of the Cas nuclease and guide RNAs into target cells, which in plants usually requires labor-intensive stable genetic transformation. However, plant viruses are emerging as a faster and more efficient alternative to deliver CRISPR-Cas reaction components into plant cells, a strategy known as VIGE^9^. Furthermore, targeting the plant germline cells with viral vectors for heritable genome editing allows to avoid plant regeneration from adult tissue^4,32^. Because novel VIGE vectors are required to broaden the host range of CRISPR, in this work we aimed to open the VIGE technology to the largest group of plant RNA viruses, genus *Potyvirus*, with most than 200 described species^33^.

A major limitation of this technology lies in the unpredictable variability of plant-virus interactions. Plant species exposed to the same virus can show dramatically different infection outcomes. This means there is substantial variability in the plant tissues the virus accesses and the resulting phenotype, observed both among different plant species and even within the same host. To overcome this, a reliable marker for specifically detecting edited plant regions was crucial. Using the CRISPOR software^27^, we designed sgRNAs to target the two homeologs of *N. benthamiana CHLI*, (**Figure 1a and S1a**)^34–36^. Of note, in the allotetraploid *N. benthamiana*, a key point for achieving a completely edited plant with a fully white phenotype is the successful editing of both alleles of each gene, which could be verified with our specific set of primers^37^.

We first compared *CHLI* editing efficiency in *N. benthamiana* plants using two different viral vectors as TRV and PVX, whose use in VIGE has already been broadly demonstrated^4,15,22,38^. Although all tested combinations of sgRNAs and viral vectors successfully induced white phenotypic patches in the infected plants, the degree of photobleaching exhibited significantly greater homogeneity when the TRV vector was employed in contrast to PVX (**Figure 1b**). This disparity is likely attributable to differences in the systemic distribution of the delivered sgRNA within the host plant. Several potential explanations for this observation exist. Firstly, TRV may possess a superior capacity for systemic colonization of *N. benthamiana* plants without imposing a substantial fitness cost or inducing severe symptoms, a characteristic well-established through its extensive application as a gold-standard vector for virus-induced gene silencing (VIGS)^39^. Secondly, the design of our sgRNAs incorporated the 5’ *FT* mobile signal, which has previously been demonstrated to enhance gene editing efficiency in systemic leaves when utilized with TRV^4^. This positive effect on RNA mobility may be different with PVX, possibly due to inherent differences in its nucleotide context or replication mechanisms that could impede the functionality of the exogenous sequence. This rationale could similarly account for variations in the sgRNA intrinsic functionality within distinct viral vector environments^25^. The ICE results directly supported our phenotypic observations, indicating that the TRV vector achieved significantly higher editing activity (**Figure 1c**). We also noted that sgRNA-1 and sgRNA-3 were generally more efficient than sgRNA-2 across both vectors. This effect was more pronounced with the PVX vector, likely because its lower efficiency as a VIGE vector allows subtle differences in sgRNA functionality to become more apparent, while a more efficient vector like TRV can mask these variations.

In a complementary analysis, we investigated the differential targeting of the two homeologous genes by the designed sgRNAs. Remarkably, we found that all sgRNAs effectively target both loci with comparable efficiency. Consequently, for future comparative studies, assessing the editing outcome of a single homeolog gene could serve as a proxy for overall functional efficacy, substantially reducing sequencing demands, at least if this reporter gene target is used. Additionally, biochemical analyses confirmed that the visible white phenotypes were indeed a consequence of altered chlorophyll synthesis. We observed an inverse relationship between chlorophyll levels and the ICE values, definitively confirming that knocking down *CHLI* leads to reduced chlorophyll accumulation (**Figure S1d**).

Following the results presented earlier, sgRNA-1 was chosen for further research, specifically to assess the inherited seed editing achieved by both vectors. In alignment with the ICE results observed in mature plants, TRV-infected plants yielded a higher proportion of edited seeds compared to those obtained with PVX. Nevertheless, the 13% edited seeds achieved with PVX, similar to what was previously reported by our group and others, serves as an encouraging demonstration, highlighting its potential utility as a vector for seed targeting in diverse host species^22,25^.

A notable observation from our germination assays was the emergence of an intermediate yellow phenotype in addition to the expected green and white seedlings (**Figure S2a**). Significantly, these yellow plants proved capable of direct cultivation on soil and subsequent seed production. Subsequent sequencing of both *CHLI* homeologs revealed a near-perfect correlation between the edited allele count and the visual color phenotype, with 30 out of 32 yellow plants presenting 3 edited alleles (**Figure 3c**). Importantly, this effect was independent of the specific *CHLI* homeolog (*CHLI-A* or *CHLI-B*) carrying the edits, underscoring the redundant and functionally equivalent nature of these two genes^37^.

Consequently, we assert that the *CHLI* marker employed in this study offers a highly practical means for identifying and growing plants on soil that possess up to 3 out of 4 edited alleles. This circumvents the necessity for *in vitro* germination of seeds carrying this genotype, a requirement for white seedlings with 4 edited alleles. We envision the development of a multiplexed system wherein an sgRNA targeting *CHLI* is co-expressed with a secondary sgRNA directed against alternative genes of interest^4,22^. Given the established positive correlation in editing outcomes between co-targeted genes in such multiplexed platforms showed in some works, yellow plants grown directly on soil could thus be readily selected and analyzed. Nevertheless, it should be taken in consideration that editing efficiency could vary substantially between different sgRNAs due to sequence context, chromatin accessibility, and other genomic factors.

After harnessing the *CHLI* visual marker to track genome editing both in adult plants and seedlings, we addressed the extension of the RNA viral vector toolbox for VIGE to the large genus *Potyvirus*^33^. They constitute the largest group of RNA plant viruses and collectively infect a wide variety of plants, including many major crops. Their collectively broad host range make them highly suitable candidates for developing versatile new VIGE vectors. Our laboratory has extensive experience utilizing potyviral-derived vectors for the expression of recombinant proteins^30,40^. Our previous research demonstrated that the intercistronic region located between the NIb and CP within the potyviral polyprotein can effectively accommodate exogenous sequences without compromising viral replication. This is contingent upon these sequences being flanked by protease cleavage sites that enable their excision from the viral proteins. Following this principle, we hypothesized that even if it did not code for a functional protein, an sgRNA could be cloned into this same position provided the open reading frame is not altered. However, the original sgRNA scaffold contains stop codons that prematurely terminate the polyprotein open reading frame in all three frames impeding production of potyviral CP. This viral protein is required for genome replication and virus movement^28^. To overcome this, we designed a modified scaffold. Building on work by Nishimasu et al.^41^, who investigated the impact of mutations on editing efficiency, we incorporated specific mutations (C25U, G46A) and deletions (ΔC30-G39) to create a novel, stop-codon-free SpCas9 scaffold (sgRNAm) (**Figure 2**). Using a TRV vector expressing sgRNA-1, we showed that the newly designed sgRNAm retained the same editing efficiency as its wild-type ancestor.

Subsequently, we proceeded to test a TEV clone incorporating the modified sgRNAm-1. Results indicated that while white patches were evident, the TEV-infected plants concurrently developed necrotic symptoms and exhibited restricted growth. Although this phenomenon confirmed the recombinant TEV vector capacity for systemic infection, it potentially exerts a detrimental effect on the overall functionality of the VIGE system (**Figure 3a**). Consistent with this visual observation, ICE analysis corroborated that TEV was capable of inducing gene editing in both *CHLI* alleles, albeit at a reduced efficiency compared to TRV. While this difference might be somewhat predictable given TEV known low protein expression levels^30^, these findings nevertheless constitute a highly encouraging initial step, underscoring the potential of this viral group for VIGE applications.

As previously discussed, a more promising and extensively researched approach for obtaining edited plants via VIGE involves heritable gene editing through germline sgRNA delivery, as it has been recently shown for the solanaceous tomato and pepper^14,15,32^. This approach facilitates the direct cultivation of edited seeds in soil, thus circumventing the need for tissue culture, a methodology that remains challenging for numerous agriculturally significant species^9^. With this objective, we decided to attenuate our TEV vector to reduce host infection symptoms and enable plants to reach the flowering stage^29^. All three attenuated TEV-sgRNAm-1 variants retained their editing activity, but only the CLA2 and AS13a variants allowed the survival and development of seeds in the infected plants (**Figure 3b-c**). As hypothesized, we observed a positive correlation between the severity of infection symptoms and the efficiency of gene editing, with the AS13a vector producing the mildest symptoms but also resulting in the least effective editing. These TEV mutants exhibit lower viral accumulation and expression of viral proteins^30^. Subsequently, we investigated the transmission of seed editing inheritance from plants infected with our attenuated TEV-sgRNAm-1 vectors. The findings indicated that 5% of seedlings derived from TEV-CLA2 infected plants possessed at least one edited allele (**Figure 4c-d**). Remarkably, RT-PCR results showed that the edited seedlings exhibited no detectable TEV infection. This outcome can be considered a valuable baseline for continued advancements in this technological platform.

Finally, we demonstrated the utility of our TEV vector for VIGE in tomato, a species of significant agronomic interest^23^ (**Figure 5a**). While the obtained ICE values were significantly lower than those observed in *N. benthamiana*, these results are a promising first step, paving the way for further exploration of this system in Cas9-expressing tomato plants. This further research must also improve heritability in crop plants, since this is a key promise of the VIGE technology. Furthermore, we extended the utility of this new VIGE system by replicating the same strategy with other potyviruses, namely TuMV and LMV, whose primary hosts are brassicas and lettuce, respectively. As we currently lack SpCas9-expressing cultivars of these agronomic species, we demonstrated their functionality in *N. benthamiana* plants. Of note, the TuMV-JPN1 strain we used in this assay is attenuated compared to the widely used UK1 strain, somehow explaining its lower capacity of systemically expressing the sgRNA^42^.

Given the conservation of biological properties within the genus *Potyvirus*, we believe these findings can be directly applied to other potyviruses, thereby broadening the range of species amenable to this technology. Nevertheless, a critical bottleneck persists within this methodology: the prerequisite for an initial round of tissue culture to establish the Cas9-expressing plant line^9^. Fully unlocking VIGE potential will therefore require all the components of the gene editing reaction must be delivered by the virus itself. Interestingly, some recent papers have reported achieving precisely this goal, employing distinct strategies across three different plant species, which demonstrates VIGE’s expanding potential^9,11,13,43^. Future attempts will be directed towards achieving the expression of novel miniature Cas nucleases in the same potyviral vectors, with the aim of obtain DNA-free non-transgenic edited seeds directly from wild-type inoculated plants^38,44,45^. Ultimately, VIGE represents a significant advancement in plant biotechnology, accelerating the pace of biological discovery and offering a powerful, accessible, and potentially regulatory-friendly platform for precise genome engineering in a wide range of economically important crops and model plants.

## Experimental procedures

### Design of sgRNAs and vector construction

The *CHLI-A* and *CHLI-B* genes from *N. benthamiana* were selected as targets for CRISPR-Cas9-mediated gene editing. sgRNAs were designed using the CRISPR-P online tool (http://crispor.tefor.net/)^46^. Briefly, sgRNAs targeting exon 3 of both genes were chosen based on high predicted efficiency and minimal off-target effects.

Plasmid pLX-TRV2 (GenBank accession number OM372496) contains an engineered TRV2 cDNA, assembled between the C*auliflower mosaic virus* (CaMV) 35S promoter and terminator, and including a hepatitis delta virus-derived ribozyme at the 3’ end^47^. pLX-TRV2 incorporates the pea early browning virus (PEBV) CP promoter ahead of a cloning polylinker with two BsaI sites (**Figure S6**). The cDNAs corresponding to sgRNA-1, 2 and 3 fused to a truncated *A. thaliana FT* sequence fragment^4^, were obtained by PCR using the high-fidelity Phusion DNA polymerase (Thermo Scientific), plasmid pPVX::sgXT2B-tFT^22^ as the DNA template, and the primer pairs D4900-D3984, D4901-D3984 and D4902-D3984 (**Table S1**), respectively. PCR consisted of an initial denaturation step at 98°C for 30 s, followed by 30 cycles of 10 s at 98°C, 30 s at 55°C and 30 s at 72°C. A final extension step of 10 min at 72°C was also included. PCR products gel separated in 1% agarose gels that were stained with 0.5 µg/ml ethidium bromide. DNA products were purified from the agarose gel using silica-gel columns (Zymoclean gel DNA recovery kit, Zymo Research). Gel-purified DNA fragments were inserted into BsaI-linearized pLX-TRV2 by Gibson assembly reactions using the GeneArt Gibson assembly HiFi master mix (Invitrogen) to produce plasmids pTRV2-sgRNA-1, -2 and 3. pTRV2-sgRNAm-1 was obtained by PCR amplifying the entire plasmid pTRV2-sgRNA-1 using Platinum SuperFi II DNA polymerase (Invitrogen) with primers D5300-D5301. This PCR reaction for long product included an initial denaturation step of 30 s at 98°C, followed by 30 cycles of 10 s at 98°C, 10 s at 60°C and 3 min at 72°C. A final extension step of 5 min at 72°C was also included. PCR products s were separated by electrophoresis in a 1% agarose gel and eluted as explained above. The gel purified full-length PCR product was circularized using the Gibson assembly reaction, as indicated above. The sequences of all plasmids generated in this work were confirmed experimentally using the Oxford Nanopore technology (Plasmidsaurus).

Plasmid pLBPVX contains a full-length PVX cDNA (GenBank accession number MT799816), flanked by the CaMV 35S promoter and terminator^48^. pLBPVXBa-M (**Fig. S6**) is a derivative of pLBPVX that contains a MluI cloning site after the PVX CP promoter, while the PVX CP itself is expressed from a heterologous promoter derived from B*amboo mosaic virus* (BaMV). The initial 29 codons of the PVX CP are deleted^49^. cDNAs corresponding to the sgRNA-1, 2 and 3 fused to the *A. thaliana* 5’ *FT* fragment^4^ were obtained by PCR as explained above for the TRV vector, but with the primer pairs D4903-D4468, D4904-D4468 and D4905-D4468 (**Table S1**), respectively. Gel-purified DNA fragments were inserted into MluI-linearized pLBPVXBa-M by Gibson assembly to produce plasmids pPVX-sgRNA-1, -2 and -3.

Plasmid pGTEVa^30^ contains the cDNA of an infectious TEV sequence variant (GenBank accession number DQ986288, with mutations G273A, A1119G), flanked by the CaMV 35S promoter and terminator within a binary vector derived from pCLEAN-G181^50^. pGTEV-G2 (**Fig. S6**) is a pGTEVa derivative that, between the NIb and CP cistrons, contains a MluI-Kpn2I polylinker flanked with sequences that complement the split NIb/CP NIaPro proteolytic site. A cDNA corresponding to sgRNAm-1 was amplified by PCR as indicated above from pTRV2-sgRNAm-1 with primers D5308 and D5309 and cloned by Gibson assembly in a MluI-Kpn2I-linearized pGTEV-G2 to produce pTEV-sgRNAm-1. Derivatives of pTEV-sgRNAm-1 containing CLA2, AS13a and AS20a debilitating mutations in the HC-Pro cistron were obtained by PCR with primers D5561-D5562, D5563-D5564 and D6077-D6078, respectively, and ligation (T4 DNA ligase, Thermo Scientific) of the products. All the primers described are listed in Table S1.

pGLMV and pGTuMV-JPN1 derive from pCLEAN-G181 (GenBank accession number EU186083) and contain a LMV sequence variant (recombinant among X97705.1 and AJ278854.1) and the TuMV JPN1 sequence variant (KM094174.2) under the control of CaMV 35S promoter and terminator. pGLMV-X and pGTuMV-K (**Fig. S6**) are derivatives with a XmaI and KasI cloning sites flanked by sequences that complement the split NIb-CP NIaPro cleavage site. cDNAs corresponding to sgRNAm-1 were amplified by PCR using the high-fidelity Phusion DNA polymerase with primers D5995-D5570 (LMV) or D5822-D5823 (TuMV) and the pTRV-sgRNAm-1 as a template, and inserted via the Gibson assembly reaction in XmaI-digested pGLMV-X and KasI-digested pGTuMV-K, respectively. Resulting plasmids were named pGLMV-sgRNAm-1 and pGTuMV-sgRNAm-1.

The complete sequence of all final plasmids used in this work is presented in **Figure S1**.

### Plant growth conditions and inoculation

*N. benthamiana* and tomato (cv. Micro-Tom) plants were grown at 25°C under a 16-hour day/8-hour night cycle. Fully expanded upper leaves from 4-to 6-week-old plants were used for inoculation with viral vectors carrying sgRNA constructs. *N. benthamiana* plants constitutively expressed a codon-optimized version of SpCas9 fused to a nuclear localization signal at the carboxyl terminus under the control of CaMV 35S promoter and *A. tumefaciens Nopaline synthase* terminator^22^. Tomato plants expressed the same SpCas9 construct, but including a DsRed reporter gene. This transgenic line was obtained by *A. tumefaciens*-mediated stable transformation as previously described^23^.

*A. tumefaciens* C58C1 electrocompetent cells carrying the pLX-TRV1 plasmid^47^ were electroporated with the pLX-TRV2 derivatives and transformed clones were selected on plates supplemented with 50 µg/ml rifampicin, 50 µg/ml kanamycin and 20 µg/ml gentamicin. For the PVX clones, the *A. tumefaciens* C58C1 strain was electroporated and transformed bacteria were selected on plates with 50 µg/ml rifampicin and 50 µg/ml kanamycin. The same C58C1 strain, carrying the pCLEAN-S48 helper plasmid^50^, was transformed with plasmids containing the different TEV, LMV and TuMV clones. In this case, transformed bacteria were selected on plates with 50 µg/ml rifampicin, 50 µg/ml kanamycin, and 7.5 µg/ml tetracycline.

Single colonies were grown for approximately 24 h at 28°C in 10 ml Luria-Bertani broth containing 50 µg/ml kanamycin and 20 µg/ml gentamicin for TRV clones, and exclusively 50 µg/ml kanamycin for the PVX, TEV, LMV and TuMV clones. At an optical density at 600 nm (OD600) ranging between 1 and 2, cells were pelleted by centrifuging at 7200 x g for 5 min and resuspended to an OD600 of 0.5 in infiltration buffer (10 mM MES-NaOH, 10 mM MgCl_2_, and 150 µM acetosyringone, pH 5.6)^51^. Resuspended bacteria were incubated at 28°C for 2 h. Two leaves per plant were infiltrated on the abaxial side using a needleless 1-ml syringe. Immediately following agroinfiltration, plants were watered. Samples from symptomatic upper leaves were collected with a 1.2-cm diameter cork borer (approximately 50 mg of tissue), immediately frozen in liquid nitrogen, and stored at -80°C until use.

### Analysis of Cas9-mediated gene editing

Leaf samples were ground with a ball mill (Star-Beater, VWR) for 1 min at 30 s⁻¹ with 1 ml of extraction buffer (4 M guanidinium thiocyanate, 0.1 M sodium acetate, 10 mM EDTA, and 0.1 M 2-mercaptoethanol, pH 5.5), and centrifuged at 15,000 x g for 5 min. Then, 0.7 ml of the clarified supernatant was loaded onto silica-gel columns (Zymo Research). After washing twice with 0.5 ml of 70% ethanol, 10 mM sodic acetate, pH 5.5, DNA was eluted in 10 µl of 20 mM Tris-HCl, pH 8.5. *N. benthamiana CHLI-A* and *CHLI-B* target fragments were individually amplified by PCR using high-fidelity Phusion DNA polymerase, as explained above and the primer pairs D5008-D5009 and D5010-D5011 (Table S1), respectively. Tomato *CHLI* target fragments were amplified by PCR using primers D5518 and D5519. *DsRED* target fragments were amplified by PCR using primers D4387 and D4388. PCR products were separated by agarose gel (1%) electrophoresis, eluted from the gel, and subjected to Sanger sequencing. The presence of sequence modifications was analyzed based on the ICE using the EditCo Bio (https://ice.editco.bio/) software.

### RNA purification and analysis of virus infectivity

Total RNA was purified from leaf tissue aliquots using silica-gel columns (Zymo Research) following the same protocol as for DNA extraction, with an added step: before loading onto the column, the collected supernatants were mixed with 0.65 volumes of 96% ethanol and centrifuged at 15,000 x g for 1 min. After purification, RNA aliquots were subjected to reverse transcription (RT) with RevertAid reverse transcriptase (Thermo Scientific). For TEV diagnosis, aliquots of the RNA preparations were subjected to RT using primer D2231, while PCR amplification was performed with primers D3599 and D3600 using the high-fidelity Phusion DNA polymerase, in the conditions indicated above. PCR products were separated via electrophoresis in 1% agarose gels, which were subsequently stained with ethidium bromide.

### Chlorophyll-a quantification

Samples collected from upper symptomatic leaves (approximately 100 mg) were mixed with 10 volumes (approximately 1 ml) of aqueous 80% (v/v) acetone precooled to -20°C, homogenized using the ball mill for 5 min at 30 s⁻¹, and mixed (1 h, 250 revolutions per min, rpm) at 4°C in the dark. Samples were centrifuged at 14,000 × g (5 min) to remove cell debris; supernatants were collected and kept in the dark until analysis. Spectrophotometric quantification of chlorophyll a (Chl-a) was performed as reported^52^. Briefly, supernatant absorbance was measured at 646 nm (A646) and 663 nm (A663) against the aqueous 80% (v/v) acetone blank in a plate reader (Infinite M Plex, Tecan), and Chl-a was quantified using the equation: Chl-a=12.21×A663−2.81×A646.

### *In vitro* regeneration of edited *N. benthamiana* plants

Upper leaves showing symptoms of viral infection were collected and submerged in water for 30 min. After that, leaves were surface-sterilized by immersion in a 50% commercial bleach solution for 10 min. Leaves were then washed three times in sterile water. Sterilized leaves were cut into pieces of approximately 1 cm^2^ using a scalpel, and were transferred to an organogenesis media (4.3 g/l Murashige and Skoog with vitamins, 30 g/l sucrose, 8 g/l bacteriological agar, 1 mg/l of 6-benzylaminopurine (BAP), and 0.1 mg/l 1-naphthaleneacetic acid (NAA), pH 5.8). Leaf discs were transferred to fresh plates every 2 weeks until shoots emerged. Shoots were cut and transferred to shoot elongation media (4.3 g/l Murashige and Skoog with vitamins, 30 g/l sucrose, 8 g/l bacteriological agar, 0.1 mg/l BAP, and 0.1 mg/l NAA, pH 5.8). Once shoots had elongated, they were transferred to root induction media (4.3 g/l Murashige and Skoog with vitamins, 30 g/l sucrose, 7 g/l bacteriological agar, and 0.2 mg/l NAA, pH 5.8). Regenerated plantlets with roots were analyzed for editing of the target genes.

### In vitro germination of N. benthamiana seeds

Seeds were collected from several flowers from plants infected with the same viral vector and pooled in Eppendorf tubes. Seeds were sterilized with 50% bleach, 0.1% Nonidet P-40, and washed with sterile water 3 times before sowing on plates with germination media (4.3 g/l Murashige and Skoog, 0.1 g/l MES, 20 g/l sucrose, 7.5 g/l agar). Seedlings were collected for genome extraction and ICE analysis.

## Acknowledgements

This research was supported by the Ministerio de Ciencia, Innovación y Universidades (MCIU; Spain) through the Agencia Estatal de Investigación (PID2023-146418OB-I00; MICIU/AEI/10.13039/501100011033 and ERDF, EU) and Generalitat Valenciana through program PROMETEO (CIPROM/2022/21). A.G. is the recipient of a predoctoral contract (FPU20/05477) from MICIU.

## Author contributions

F.M. and J.A.D. conceived the work with input from the rest of the authors. F.M., V.A. and A.G. performed the experiments. All authors analyzed the data. F.M. and J.A.D. wrote the manuscript with input from the rest of the authors. All authors discussed and revised the manuscript.

## Data availability statement

The data that supports the findings of this study are available in the supplementary material of this article.

## Supporting information

**Figure S1.** Nucleotide sequences of the viral vectors used in this work.

**Figure S2.** (a) The N. benthamiana magnesium chelatase participates in the metabolic pathway of chlorophyll biosynthesis. Three different guide RNAs targeting simultaneously the two homeologs of N. benthamiana Magnesium chelatase subunit I (CHLI) were designed with the CRISPOR tool (http://crispor.tefor.net/), considering their predicted efficiency and minimizing off-targets. (b) Time-lapse pictures encompassing 14 dpi showing the performance of the TRV-sgRNA-1 vector editing CHLI. (c) Representative adult plants agroinoculated with vectors TRV-sgRNA-1, -2 and -3 (top line) and PVX-sgRNA-1, -2 and -3 (bottom line), as indicated. (d) Chlorophyll a accumulation in systemic leaves of N. benthamiana plants inoculated with the different viral vectors expressing sgRNA-1, -2 and -3, as indicated. Pictures of harvested leaves are on the left. Graph (on the right) corresponds to average chlorophyll a accumulation of three different upper leaves for each treated plant. Error bars indicate standard deviation.

**Figure S3.** (a) Phenotype of the seedlings from *N. benthamiana* plants infected with TRV-sgRNA-1 could be divided in three categories: green (G), light green/yellow (Y) and white (W). (b) ICE analysis of both homologues from 32 individual seedlings of each phenotype. Individual seedlings were plotted indicating its phenotype and the number of edited alleles. (c) Yellow plants with 3 out of 4 edited alleles are able to grow on soil and produce functional flowers. (d) According with a Mendelian transmission trait, the seedlings coming from these plants presented the expected 1:2:1 (white:yellow:green) segregation.

**Figure S4.** (a) RT-PCR diagnosis of TEV in N. benthamiana plants regenerated in vitro. All plants are positive for TEV detection, except the negative control (not infected adult plant). (b) RT-PCR diagnosis of TEV in N. benthamiana seedlings. Green arrow in lane 22 shows the only positive TEV positive sample. Lines 1, 13, 19 and 24 correspond to edited seedlings in which TEV is not detected by RT-PCR.

**Figure S5.** Expanding the use of the TEV vector with the modified SpCas9 scaffold to another gene target. (a) Micro-Tom tomato expressing SpCas9 and DsRED were agroinoculated with TEV-sgRNAm-R1, targeting the DsRED gene. Images show one leaf of a non-inoculate control plant (left) and two leaves of an agroinoculated plant (center and right). b) Graph of ICE values of 11 analyzed samples of DsRED-SpCas9 tomato infected with TEV-sgRNAm-R1. A representative Sanger sequencing profile is shown. Sequence homologous to sgRNAm-R1 protospacer is boxed.

**Figure S6.** Nucleotide sequences of intermediate cloning vectors used in this work.

